# CATHI: An interactive platform for comparative genomics and homolog identification

**DOI:** 10.1101/2023.09.04.556229

**Authors:** Lukas Becker, Philipp Spohr, Gunnar W. Klau, Ilka M. Axmann, Sebastian Fraune, Nicolas M. Schmelling

## Abstract

Bioinformatics has established itself as a central pillar of modern biology. Specifically, comparative genomics enables scientists to study a vast number of genomes efficiently. These comparative analyses shed light on the evolution and potential function of genomes and genes, but are also increasingly used as a key tool for metabolic engineering and synthetic biology by identifying appropriate targets for modification. While numerous sophisticated tools for comparative genomics and homolog identification exist, those tools predominantly target highly skilled bioinformatics users. Consequently, many biologists either defer such analyses to their more versed bioinformatic collaborators or resort to suboptimal tools. Here, we present an intuitive solution available on all major operating systems, easily accessed through common web browsers. CATHI – Comparative Analysis Tool for Homolog Identification – integrates a suite of best-practice bioinformatic tools, encompassing BLAST for homology searches, MAFFT for multiple sequence alignment, FastTree2 for phylogeny reconstruction, and clinker for synteny analysis. Specifically tailored to biologists, CATHI orchestrates predefined settings and automated pipelines, obviating the need for programming expertise. This platform empowers researchers to confidently engage in detailed comparative genomics studies by streamlining the analytical process. The interactive framework provides users with a plethora of options. This includes real-time execution and progress monitoring, facilitates dynamic result tracking, and a set of search functions across NCBI databases like CDD or ProtFam. Users can interactively engage in data exploration, filtering, and visualization through CATHI’s intuitive interface. Furthermore, the seamless export of project data in standard formats (FASTA, Newick, CSV, and HTML) facilitates the integration with further third-party tools such as TreeViewer and Jalview. To benchmark CATHI, we revisited the comparative analysis of cyanobacterial circadian clock proteins conducted by Schmelling et al. in 2017, revealing consistent global patterns among identified homologs, while also highlighting individual variations attributed to the expansion of available databases.

## 1 Introduction

Bioinformatics has emerged as a critical cornerstone of modern biological research, reshaping the landscape of biological exploration and transforming it from traditional laboratory confines into a dynamic, data-driven, and multidisciplinary field. Comparative genomics allows researchers to compare and contrast genomes from different species. Those analyses provide insights into evolutionary relationships, identifying conserved genes and regions, and understanding genetic variations that underlie phenotypic diversity. A key concept of evolutionary and comparative genomics is the dichotomous differentiation of homology into orthologous and paralogous sequences [1, 2]. Paralogs are genes related by an ancestral gene duplication event, while orthologs are homologous genes in different species descended from a single ancestral gene of a last common ancestor through vertical inheritance and subsequent speciation [3]. Sets of orthologous genes are used to obtain information regarding gene function and phylogenetic relationships. Thus, a clear distinction between orthologous and paralogous genes is crucial for reliable and robust genome annotation and phylogenetic inference [2, 4]. However, those definitions were proposed before the advent of the genomic era. Assigning orthology or paralogy for genomic sequences is much more complex as sufficient information is still missing to “determine the timing of many of the speciation and gene duplication events” [5]. Thus, it is suggested to classify homologs based on sequence-structure-function relationships as “isofunctional” or “heterofunctional” and “isospecic” or “heterospecic” homologs [5].

Typically, orthologous and paralogous gene relationships are disentangled by analyzing sequence similarities and their distribution within phylogenies [1, 6]. Different computational solutions are required depending on the biological question [7, 8]. Thus, various tools have been developed to infer orthologs [9]. Some of these tools, like OrFin [10], OrthoMCL [11], and OrthoFinder [12], use a combination of existing bioinformatic software, such as BLAST (Basic Local Alignment Search Tool; [13, 14, 15]), DIAMOND [16], MMseqs2 [17], Markov Clustering (MCL) [18], or phylogenetic reconstructions. Other tools are based on different approaches to identify orthologs, like SPOCS [19], OrthoInspector [20] or JustOrthologs [21]. SPOCS combines the results of pairwise species comparisons from the InParanoid [22] algorithm with graph-based predictions of orthologs and paralogs [19], and aims to identify orthologs within species groups. OrthoInspector [20] validates the output of sequence similarity searches and creates groups of in-paralogs that are compared pairwise to assign potential orthologs and/or in-paralogs, while JustOrthologs [21] uses coding sequence lengths and dinucleotide percentages to assign orthologs. Moreover, the focus of these tools or their respective online databases are different. While morFeus [23] is specifically designed for identifying remotely conserved orthologs, SPOCS, on the other hand, is aimed at identifying orthologs within closely related species [19]. These tools differ in their installation, operating system compatibility, and usage (Table 1).

**Table 1:**
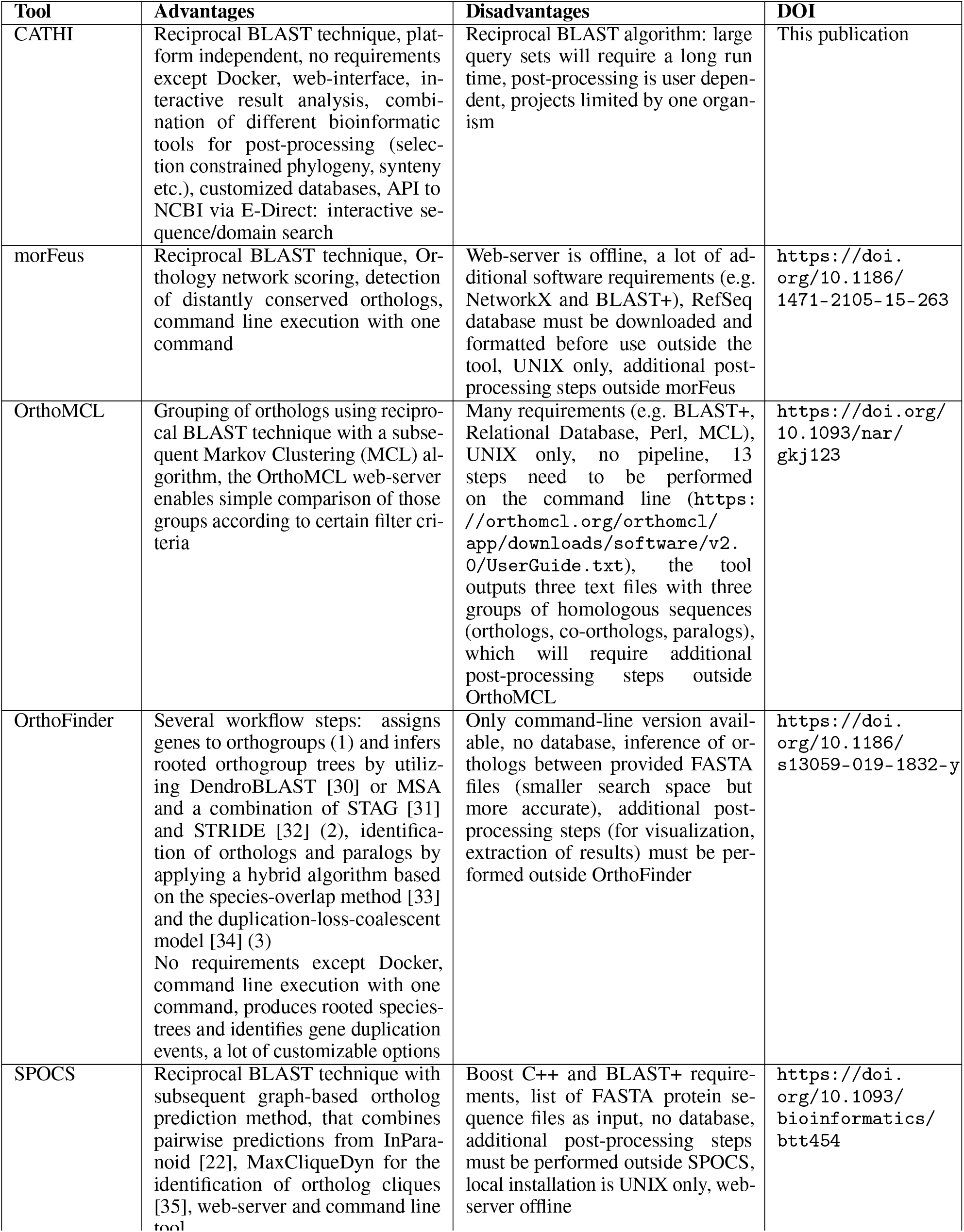
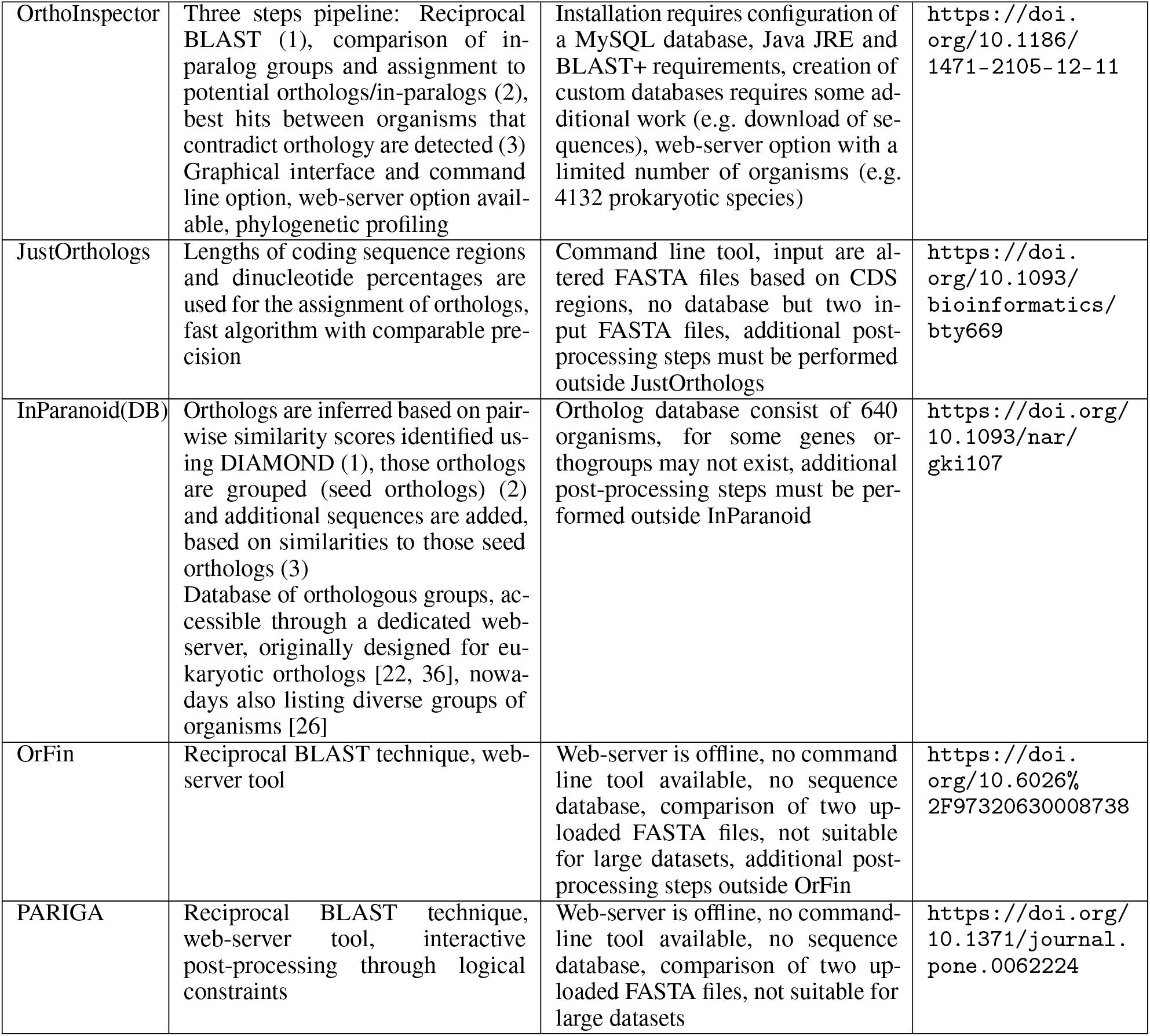
Brief comparison of a selection of analytical tools for the identification of homologs and orthologs.

In addition, various biological databases like the Cluster of Orthologous Groups (COG) database [24], OrthoDB [25] or InParanoidDB9 [26] have been developed to store groups of orthologous and/or paralogous sequences, allowing for rapid identifications of homologous relationships among genes and species. EukProt [27] and PanOCT [28] are databases that primarily focus on identifying orthologs among eukaryotic and prokaryotic genomes, respectively. In contrast, more generalized approaches like the triangular Reciprocal Best Hit (RBH) method used in the COG database [24] take a broader approach to unravel the homologous relationships among genes.

Still, in most of these tools or their respective databases, homologs are (at least partly) identified through RBHs (Table 1). RBHs can be assigned by performing a reciprocal or symmetric BLAST search (Fig. 1). BLAST rapidly searches large sequence databases by finding local similarities. Local alignments identify regions of similarity between two sequences, highlighting regions that might share a common function. BLAST starts with a short sequence, searches for sequences sharing similarities, and extends those alignments as best as possible by applying cut-off values based on sequence substitution matrices [13]. In contrast, global alignments use the complete sequences and try to best align them with one another [29].

**Figure 1.**
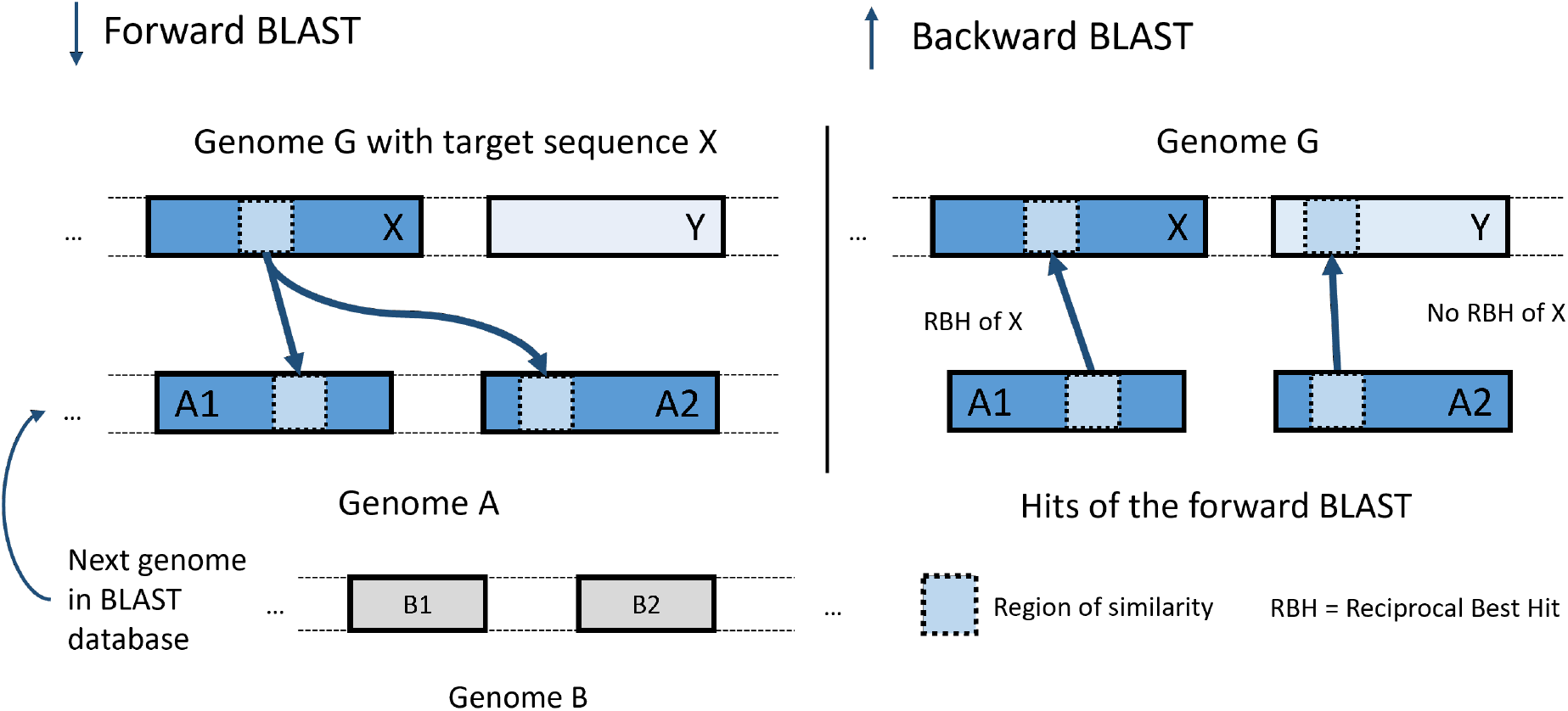
Schematic overview of reciprocal BLAST. Genome G is the target organism from which the query sequence X originates. X is the input (query sequence) for the first BLAST, the forward BLAST, left. The results of the forward BLAST (sequences A1 and A2) are the inputs to a second BLAST, the backward BLAST, right, which uses the query genome G as a database. A1 finds X as its best match, while A2 aligns best with Y. Thus, only A1 is a reciprocal best hit (RBH) to X.

Within a reciprocal BLAST approach, the sequence(s) of interest (SOI) from a specific species are used as input queries for the initial sequence similarity search, also known as the forward BLAST (Fig. 1, left). The generated output is then analyzed, and the sequences of all hits from the forward BLAST are extracted from the underlying database. These extracted sequences then serve as queries for the second sequence similarity search, also known as the backward BLAST (Fig. 1, right). In this step, the BLAST is constrained by explicitly searching against all sequences of the initial organism, also known as the query genome. Additionally, the output is limited to reporting only the best hit or alignment. RBHs are assigned if the query and subject sequences from the forward BLAST match against each other in the backward BLAST (Fig. 1, right).

Although sophisticated tools are available for identifying homologs (Table 1), to our experience, many biologists still do not use them. Those tools predominantly target bioinformaticians as their potential users. Thus, biologists leave those analyses to their bioinformatic collaboration partners or use less suitable tools, like the online BLAST tool available on NCBI, to identify homologs. While the online BLAST tool on NCBI is tremendously helpful to biologists, it offers limited options for correctly identifying homologs. Thus, it should instead only be used for the initial screening of candidates. In our experience, biologists tend to use tools that (1) have a graphical interface, (2) do not require any complicated installation process or command line experience, (3) can be used without programming skills, and (4) already offer many useful default settings. To overcome this discrepancy and also provide biologists with sophisticated tools to perform high-quality comparative genomic analyses and homolog identifications confidently, we combined a set of state-of-the-art tools and best-practice analyses (see Results - Identification of Homologous Sequences; Multiple Sequence Alignments and Phylogenetic Reconstruction; Genomic Structure and Synteny Analysis; Conserved Domain Database (CDD) Analysis) into an easy-to-use interactive tool with an intuitive graphical interface (see Results - Interactive Filtering).

Here, we introduce an accessible solution that can be effortlessly installed on all major operating systems, enabling straightforward usage through an interactive and intuitive web interface. The default settings are selected to provide a starting point for any homology search. However, more sophisticated users can change all settings to their personal preferences. Further, we illustrate CATHI’s functionality and reliability through a reproduction and benchmarking analysis of cyanobacterial circadian clock proteins based on Schmelling et al., 2017 [37]. This analysis revealed that overall trends in homology detection are consistent even with substantially increased database size. However, individual variations have been identified highlighting the need to thoroughly interpret results of comparative genomics analysis, especially when using different databases.

## 2 Results

CATHI combines a suite of powerful bioinformatic tools, including BLAST+ [15] for homology searches, MAFFT [38] for multiple sequence alignments (MSA), FastTree2 [39] for phylogeny reconstruction, and clinker [40] for synteny analyses into an interactive platform for comparative genomics specifically tailored to biologists. The predefined settings and pipeline automation streamline the process and enable researchers to perform sophisticated comparative genomics analyses regardless of their proficiency in programming or bioinformatics. The interactive and intuitive web interface provides users with multifaceted capabilities, including execution and monitoring, real-time result tracking, search function across diverse NCBI databases (Protein, CDD, ProtFam), and dynamic data exploration, filtering, visualization, and post-processing. CATHI runs on all major operating systems supporting Docker [41] and is accessible via typical browser applications (e.g., Firefox or Chrome). Through individual accounts, users have the option to create their custom projects and retrieve their comprehensive project data in standard formats (FASTA, Newick, CSV, HTML) suitable for further analysis with other third-party tools such as TreeViewer [42], Jalview [43], or vector design programs. In the following, we highlight the main features of this platform and provide examples of potential use cases, illustrated through the replication of the prior comparative study on cyanobacterial circadian clock proteins by Schmelling et al., 2017 [37].

### 2.1 Identification of Homologous Sequences

The homology search module within CATHI represents the core of each project. Users start each comparative genomics analysis by creating a new project through the web browser dashboard (Fig. 2). Before project initiation, users either choose from already formatted BLAST databases or create a new database using CATHI’s database creation module (Fig. 2). Databases can range from the entire GenBank or RefSeq database from NCBI, over an individually filtered subset of those databases, to directly uploading annotated genome files. The latter option allows users to incorporate unpublished genome projects into their custom database. By entering the SOI(s) of one organism per project, users start the reciprocal BLAST analysis with either default (see Methods - BLAST and Identification of Reciprocal Best Hits) or custom settings. The automated pipeline then performs all steps of forward and backward BLAST as well as filtering without any further input from the user (Fig. 2 -Homology Search Module). Afterward, CATHI outputs a set of analysis statistics. Furthermore, the BioPython [44] library extracts taxonomic information of the organisms listed in the RBH results table from NCBI. Subsequently, these taxonomic data are integrated into the results table and written to the project directory (Fig. 2 -Interactive Results Table). All data tables can be downloaded as CSV files or are conveniently accessible within CATHI, facilitated by integrating the client-side JavaScript library DataTables (http://datatables.net). This versatile library enriches user experience with its suite of advanced functionalities, enabling seamless searching, sorting, paging, and dynamic interaction with extensive HTML tables. Through the homology search module, users can perform reliable reciprocal BLAST analysis with just a few clicks and inputs (Fig. 2).

**Figure 2.**
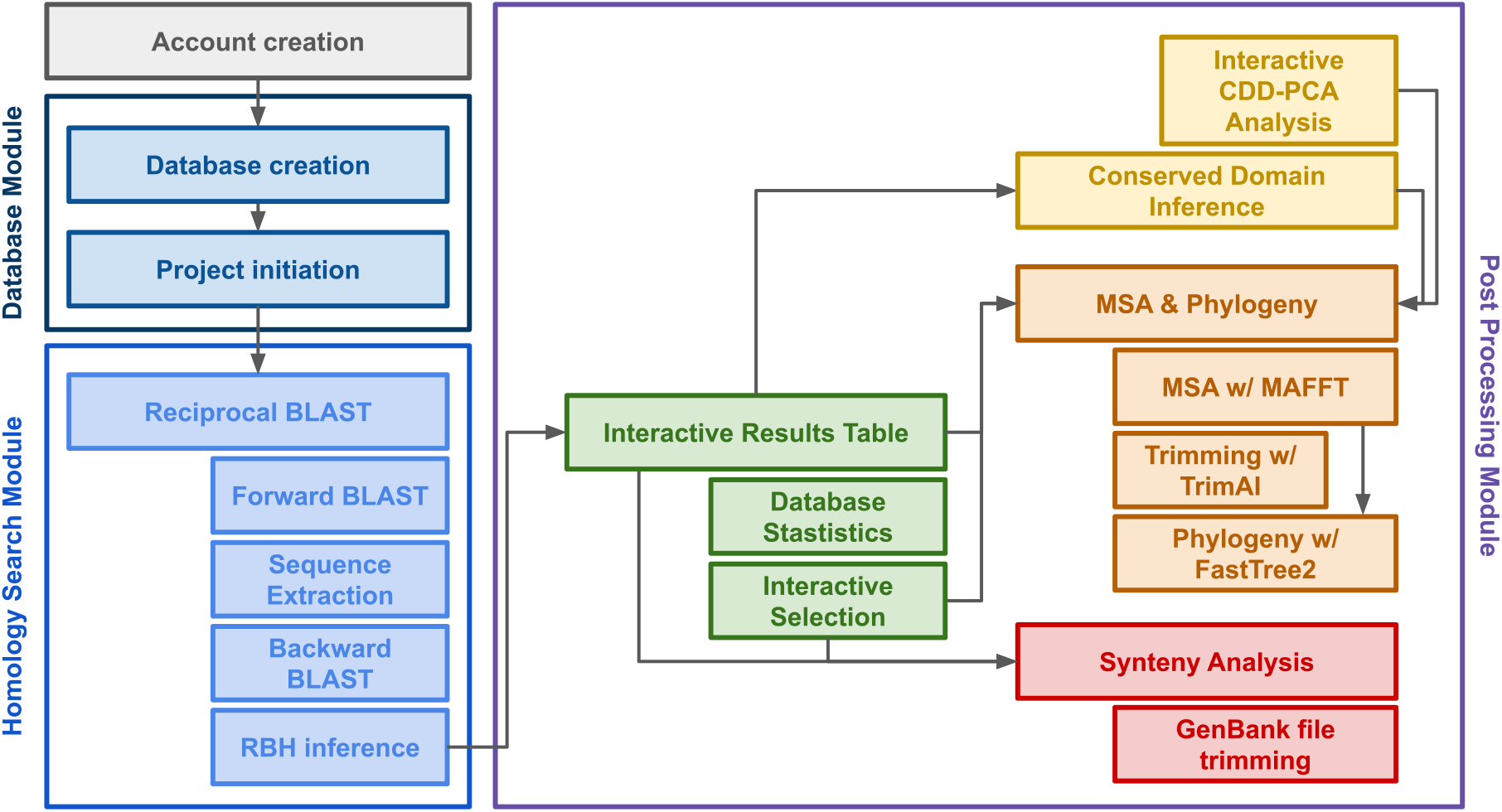
Flowchart of CATHI’s pipeline, modules, and post-processing options. First, users have to create an account to login to the platform. They can use the **Database Module** to create a database from locally uploaded protein FASTA files or remotely available proteomes of the RefSeq or GenBank. Users can initiate a new project by selecting or creating appropriate databases, providing SOIs from one organism of interest per project and optionally customizing the default settings. After database and project creation, users perform the reciprocal BLAST analysis using the **Homology Search Module**. The underlying pipeline within the module will then automatically perform the forward and backward BLAST, sequence extraction, and reciprocal best hit (RBH) inference. The output of this module is directly forwarded to the **Post Processing Module**, where the users have different options to interactively analyze and filter their RBH data. Within the Post Processing Module, interactive results tables and statistics are available, synteny analyses and conserved domain interference can be performed, as well as multiple sequence alignments (MSA) and phylogenetic reconstruction. Furthermore, those different analyses can be combined into interactive pipelines in which users can first filter their RBH data by interactive graphs and tables before constructing MSAs and phylogenies.

### 2.2 Multiple Sequence Alignments and Phylogenetic Reconstruction

To identify conserved regions and patterns of similarity/difference within the identified RBHs and to elucidate the evolutionary relationship between them, post-processing involves the construction of an MSA and phylogenetic tree for each set of RBHs (Fig. 2 -MSA & Phylogeny). MAFFT (Multiple Alignment using Fast Fourier Transform) is used to conduct the MSA ([38]; Fig. 2). Its primary purpose is to align multiple sequences with homologous regions to identify conserved regions and evolutionary relationships, which makes it perfectly suitable for CATHI. Subsequently, the alignment undergoes a refining process using trimAl [45], a specialized tool that automates the removal of inadequately aligned segments (Fig. 2). The trimming not only enhances the precision of the alignment but also contributes to the refinement of phylogenetic inference, ensuring a more robust and accurate analysis. To highlight conserved regions within the MSA and create an interface that allows interactive visualization, an HTML document is created using the MView [46] tool. Phylogenetic inference is performed with FastTree2 ([39]; Fig. 2). It leverages sophisticated approaches to approximate maximum-likelihood phylogenies. Subsequently, resulting phylogenetic trees are processed by shiptv (https://github.com/peterk87/shiptv) to generate interactive HTML visualizations (Fig. 3). This visualization integrates the RBH results table by aligning the data to the phylogenetic tree leaf labels, facilitating a comprehensive and insightful depiction of sequence relationships. Through the integration of the data tables, users can interactively filter and adjust the phylogenetic tree based on selection criteria such as taxonomic groups (Fig. 3).

**Figure 3.**
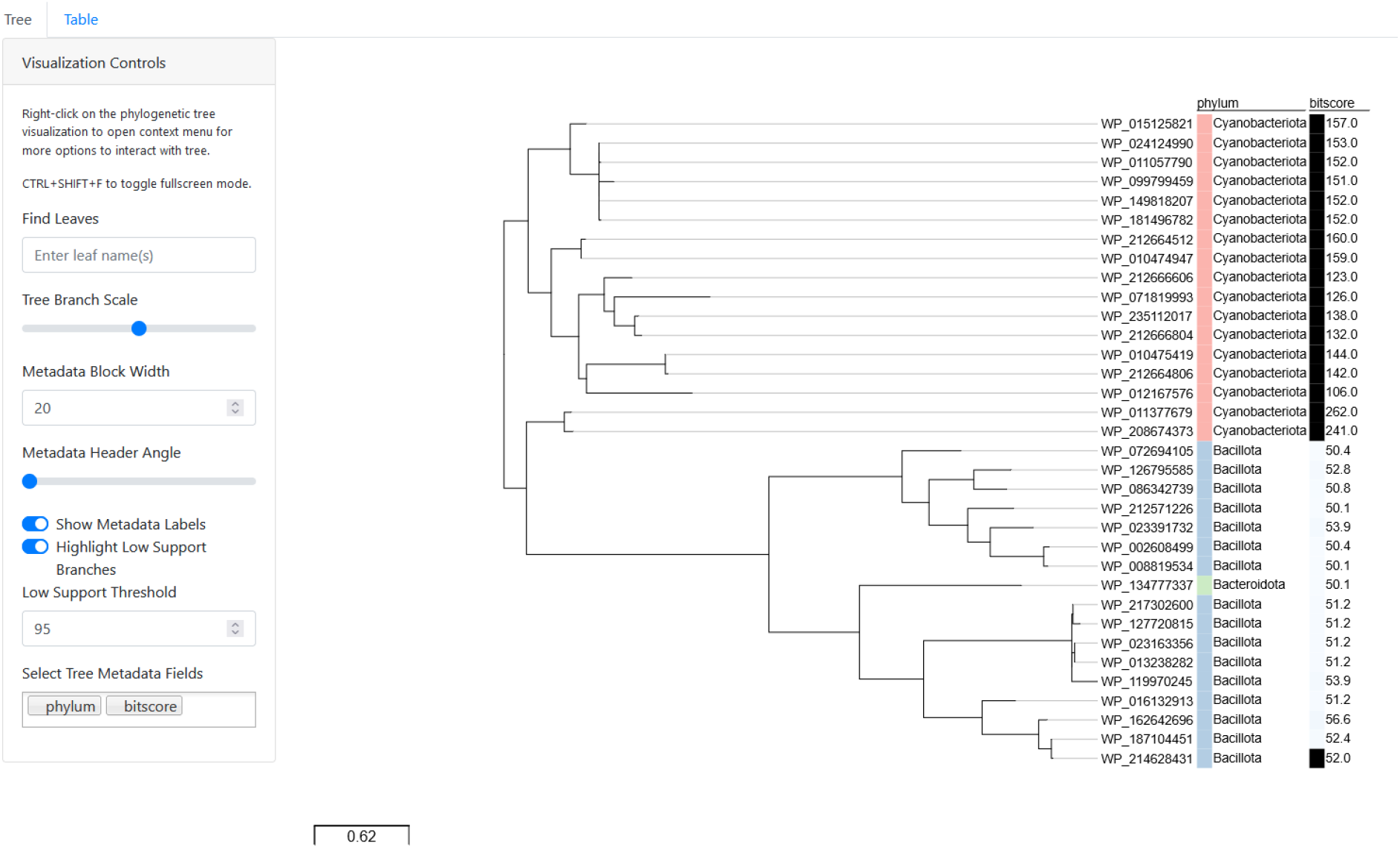
Interactive Phylogeny Visualization. The interactive phylogeny visualization is presented here as an exemplary illustration. This dynamic visualization was generated using a set of bioinformatic tools: MAFFT provides accurate multiple sequence alignments, FastTree2 enables rapid and efficient phylogeny reconstruction, and shiptv facilitates the creation of an interactive dashboard view. The tree’s branches represent evolutionary relationships, while the distribution of a selection of RBHs of the cyanobacterial circadian clock input factor Pex is highlighted across three bacterial clades, clearly distinguishing between *Cyanobacteria* and *Bacillota*.

### 2.3 Genomic Structure and Synteny Analysis

Beyond RBHs identification, MSA, and phylogenetic reconstruction, users can analyze the genomic structure and synteny surrounding the RBHs (Fig. 2-Synteny). Therefore, CATHI integrates the clinker tool, which allows researchers to compare and visualize syntenic regions across different genomes [40]. To perform this analysis, users can select up to ten RBHs of interest, and CATHI will download the corresponding GenBank files from NCBI and extract gene loci around each RBH. These sliced GenBank files are automatically passed to clinker as input, enabling users to easily identify and analyze conserved gene clusters and gene order between genomes directly from their RBH data (Fig. 4).

**Figure 4.**
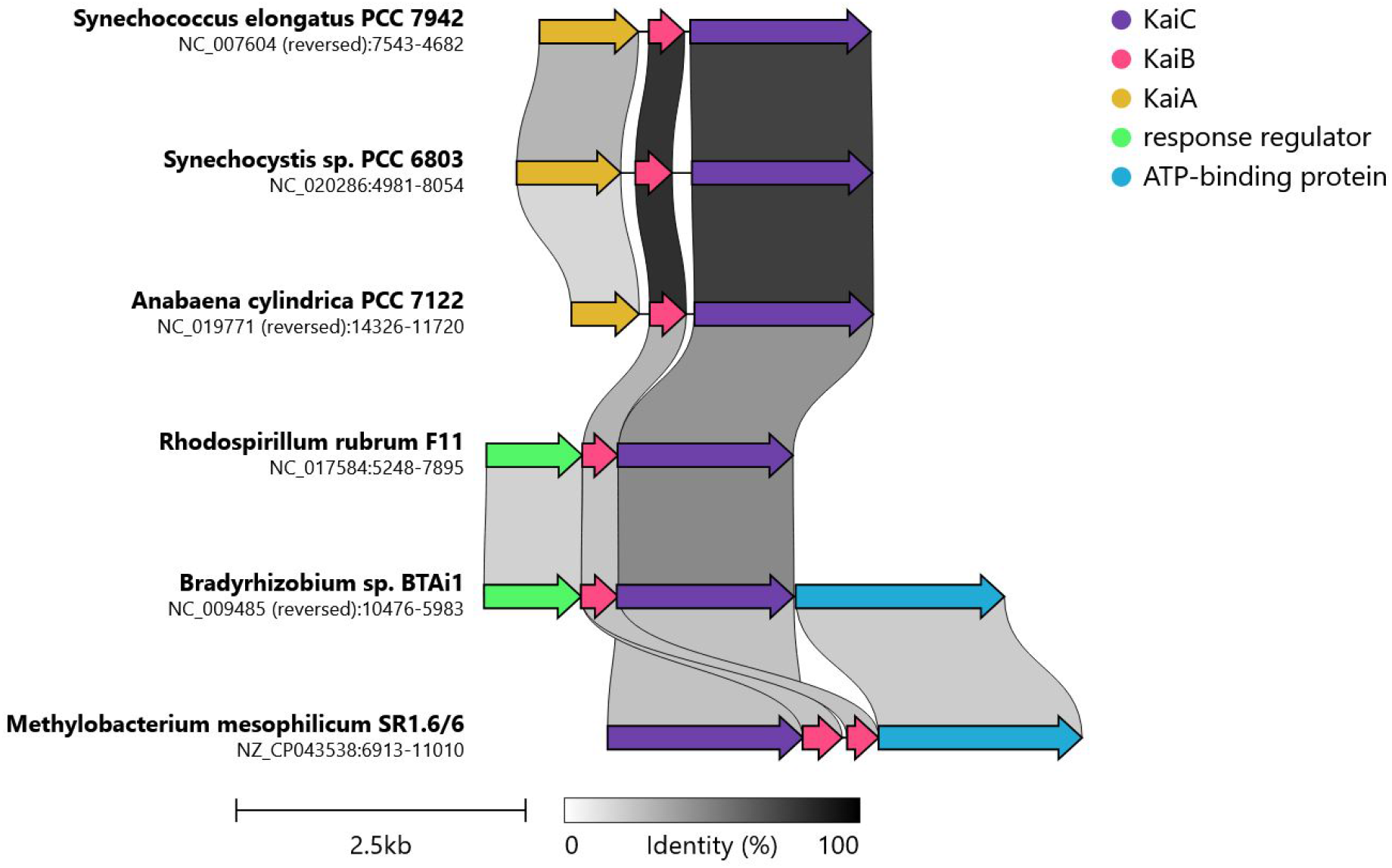
Exemplary illustration of the synteny analysis results using clinker as part of CATHI’s post-processing module. In this analysis, the core circadian clock protein KaiC was selected. Clinker performed the synteny analysis. The graph shows the conserved genomic elements up-and downstream of KaiC that are represented in the selected genomes. The threshold for sequence identity is set to 0.25 (25%). Interestingly, the synteny analysis reveals some intricate information about the KaiABC cluster without any additional input that can easily be missed in standard RBH analyses. First, the *Anabaena/Nostoc* KaiA has a smaller sequence length compared to the KaiA sequences of *Synechococcus elongatus* PCC 7942 and *Synechocystis* sp. PCC 6803. Additionally, the synteny reveals the presence of a conserved response regulator/transcription factor in the genomes of the *Bradyrhizobium* and *Rhodospirillium rubrum* strains at a position where normally KaiA can be found. This protein was recently identified as a novel KaiA homolog and was named KaiA3 due to its functional similarity [47].

### 2.4 Interactive Filtering

CATHI offers a set of additional features to analyze the RBHs based on taxonomic information and database entries (Fig. 2 -Interactive Selection). The number of available genomes of an organism (e.g., *E. coli*) in the BLAST database may vary, affecting the analysis of RBHs due to high data redundancy. Therefore, project-specific database statistics are calculated based on the taxonomic information within the RBH results tables and the underlying BLAST database. Each RBH is derived from a genome file of one organism. CATHI calculates database-normalized percentages of taxonomic units within identified RBHs. The resulting database statistics are integrated into the RBH results table and visualized by an interactive chart that combines RBH characteristics and a table of database entries. The table and the chart are linked, and the table data is updated according to the specified user selection (Fig. 5). This comprehensive graph empowers users to filter RBHs based on various features, including bitscores, RBH sequence lengths, sequence identity, e-value, and taxonomic nodes (Fig. 5). To facilitate user interaction, a lasso tool is provided, enabling the direct selection of RBHs within the graph (Fig. 5). The database entry table is thereby filtered by the selected RBHs. The selected RBHs can be downloaded or utilized for the creation of multiple sequence alignments and phylogenetic inferences (Fig. 2). By leveraging this interactive graph, users can effectively explore and analyze the RBHs, gaining valuable insights and facilitating the identification of relevant sequences. The integration of RBH features and database statistics within the visual representation enhances the usability and interpretability of the results. The number of representative proteins can be examined and compared in other organisms and across phylae, classes, orders, families, and genera.

**Figure 5.**
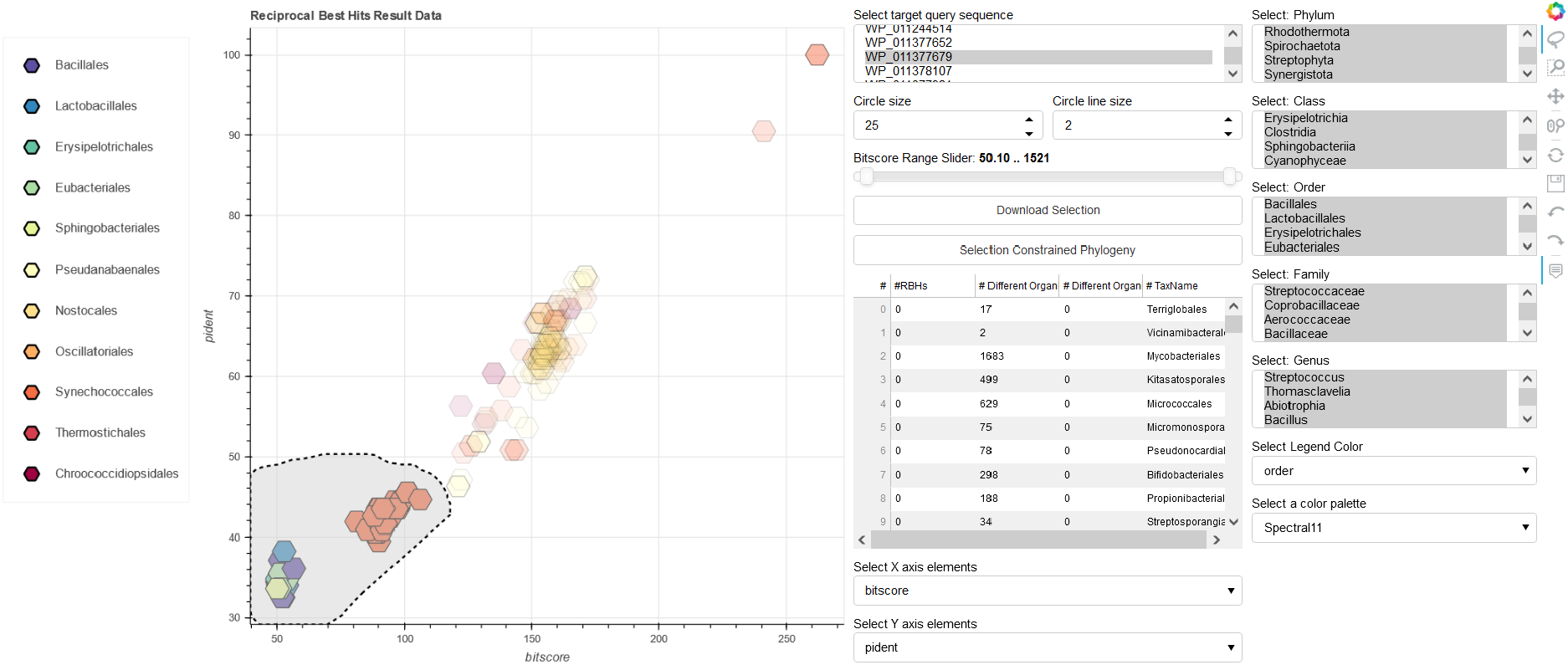
Interactive reciprocal best hits analysis and filtering. The interactive scatter plot leverages the reciprocal best hit (RBH) results dataset, exemplified here with the Pex protein dataset. The X and Y axes of the scatter plot are parameters of the dataset, such as bitscore, e-value, sequence identity (pident), and sequence lengths, enabling the exploration of relationships among these RBHs. The interactivity of the scatter plot is facilitated by Bokeh, a powerful visualization library [48]. Users can dynamically manipulate the dataset through filtering options informed by the taxonomic information associated with each RBH. In addition, Bokeh provides a lasso tool to select specific RBHs within the graph. The lasso tool can be selected from the Bokeh tool-panel displayed at the right side of the figure. This interactive feature empowers researchers to dissect real-time taxonomic trends and relationships among RBHs, unveiling underlying patterns and insights. In this example featuring the Pex protein dataset, the scatter plot offers a multidimensional view of the RBHs landscape. As users navigate through the data points, taxonomic affiliations emerge, shedding light on the distribution and relationships of Pex protein homologs across various taxa. The provided interactivity not only enhances the visualization experience but also grants researchers the capability to refine their analyses, making the scatter plot an indispensable tool for exploring the taxonomic diversity of RBHs.

### 2.5 Conserved Domain Database Analysis

To gain further knowledge about SOI(s) CATHI offers the possibility to infer protein domains for the SOI and the associated RBHs (Fig. 2 -Conserved Domain Interference). The domains are inferred from a local copy of NCBI’s Conserved Domain Database (CDD) [49]. CATHI generates result tables of protein domains that are transferred into interactive HTML tables for visualization and analysis. Additionally, CATHI writes a FASTA file containing the concatenated domain sequences of the RBHs. This FASTA file is then used for an additional MSA and refined phylogenetic reconstruction, as highlighted before. Furthermore, a principal component analysis (PCA) is conducted on a percent identity (RBH vs. CDD entry) table of the conserved domains (Fig. 2). A graph of the first two principal components of the PCA in combination with a domain table is visualized in an interactive chart, with the possibility to filter RBHs based on taxonomy, similar to the interactive database statistic graph (Fig. 6). The selected RBHs can be downloaded for studies outside CATHI or used for a selection constrained phylogenetic inference based on the identified protein domains (Fig. 2).

**Figure 6.**
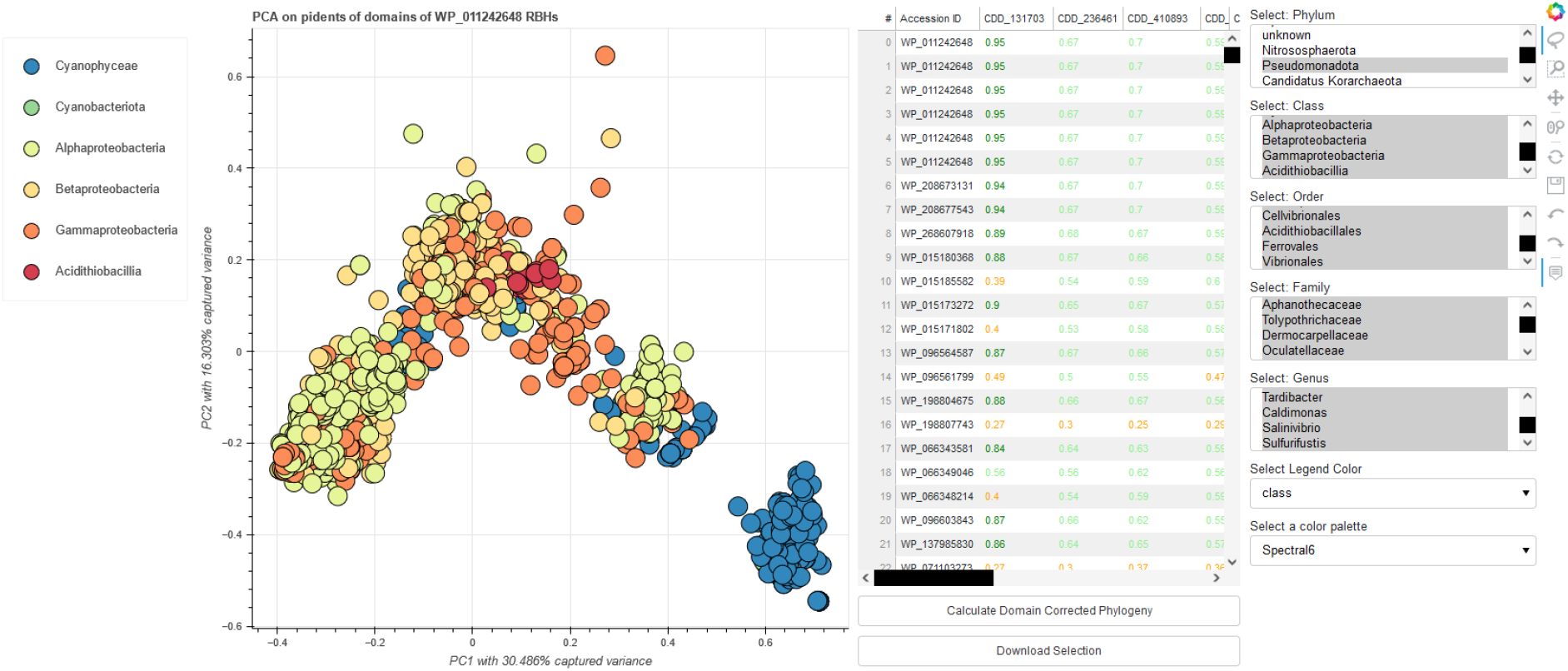
Interactive graph of inferred CDDs of *Synechococcus* KaiC RBHs. The figure displays the first two principal components of a principal component analysis (PCA) calculated on a percent identity table (RBH vs. CDD entry) of protein domains (displayed in the table next to the graph) inferred from RBHs of the *Synechococcus* KaiC protein. The graph and table are linked, selection of specific scatter-points will alter the percent identity table. Coloring within the graph is based on the associated taxonomic class, coloring within the table is based on the percent identity value. The figure displays a selection of organisms from the *Cyanobacteria* and *Pseudomonadota* phylum. Some of the RBHs are known KaiC2/C3 cyanobacterial proteins, paralogs of the query sequence KaiC. Interestingly, the cyanobacterial RBHs most similar to the *bona fide* KaiC form a cluster in the lower right corner of the diagram. They exhibit differences in their domain structure compared to the other RBHs.

### 2.6 Reproducibility

The amount of biological data has increased dramatically and will continue to grow, rendering data comparisons and analysis more complicated. Consequently, scalable software tools and reproducible bioinformatic analysis are essential for modern comparative genomics [50]. Within CATHI we implemented a set of measures to ensure and enhance the reproducibility and integrity of each analysis. First, deploying individualized local accounts prevents the accidental merger of projects and ensures a clear dashboard for each user. The platform enforces unique project names by design to prevent inadvertent data overwrite. Further, each BLAST database is created as a discrete and unmodifiable database, thereby preserving the content of the database at the time of initiation and insulating it from subsequent updates. This approach allows users to re-execute analyses at any time, building a dynamic environment that promotes the iterative validation and refinement of results. CATHI’s architecture is grounded in utilizing a PostgreSQL database equipped with the inherent advantage of data persistence. This resilience against local file deletion safeguards against inadvertent data loss, ensuring the persistence of each dataset. Through containerization with Docker, the specific version of the underlying tools is preserved and recorded, simplifying the reporting of software versions. Furthermore, the orchestration with Snakemake [51] amplifies the automation of the pipeline and ensures the correct execution, reducing potential errors from manual data manipulation and execution of tools. To further increase reproducibility, Snakemake configuration files are written into each underlying project folder, and corresponding Snakefiles and scripts can even be used outside the tool.

Each created database and analysis is assigned a timestamp. In addition, log files are generated, recording the exact steps and execution of each analysis. The resulting raw and processed data of each analysis are preserved in their entirety, while any filtering or transformation generates discrete datasets that augment interactive exploration and evaluation. This approach circumvents the overwriting of original data, encouraging users to explore and manipulate data without fear of irrevocable data loss. Incorporating interactive HTML tables allows each data table to be accessed through the platform. Further, each analysis result can be downloaded in a standardized file format, such as CSV for data tables, FASTA for sequence and alignment files, and Newick for phylogenetic trees. Conforming to standardized file formats further enhances the reproducibility of findings across disparate computational pipelines, ensuring interoperability and knowledge exchange.

### 2.7 Benchmarking

To assess the performance of CATHI’s reciprocal BLAST pipeline, we conducted a comparative analysis of circadian clock proteins, as previously described in Schmelling et al., 2017, Wiegard et al., 2013 and Axmann et al., 2014 [37, 52, 53]. The cyanobacterial circadian clock primarily consists of three key proteins: KaiA, KaiB, and KaiC. KaiC, in particular, plays a crucial role and exhibits ATPase, auto-phosphatase, and auto-kinase activities. Activation of KaiC occurs through KaiA-mediated auto-phosphorylation at specific amino acid residues, while KaiB acts as an antagonist by occupying potential KaiA binding sites [54].

We performed the analysis on the same set of protein sequences used in the study by Schmelling et al., 2017 [37] (Table S1). To achieve this, we utilized CATHI’s database features to download and format RefSeq proteomes into BLAST databases. The entries were filtered automatically by CATHI based on the completeness levels “Complete Genome” and “Chromosome,” resulting in a final database containing 53,657 proteome entries with 28,326 unique taxonomic identifiers from 94 different phylae and 168,314,551 protein sequences. Within this database, there are 280 proteomes from 265 different cyanobacterial species. The database used in the analysis conducted by Schmelling et al., 2017 comprised 11,300 proteomes, including 94 cyanobacterial proteomes from 64 distinct cyanobacterial organisms. Each of these proteomes was used as a single BLAST database, which minimizes the e-value for possible homologous sequences, whereas CATHI uses a single BLAST database consisting of all downloaded proteomes. Following CATHI’s best practices, we created two reciprocal BLAST projects: one for 16 *Synechococcus* query proteins and another for six *Synechocystis* query proteins (Table S1). The proteins WP_011378436 and WP_041425845 used in Schmelling et al., 2017 have been suppressed from NCBI as they are no longer annotated on any genome and are excluded from this study. Instead of the suppressed WP_041425845, the updated version under the accession number WP_010873802 was used, while WP_011378436 was excluded from the analysis. All remaining query proteins were extracted from the result directories provided in Schmelling et al., 2017, and removed sequence identifiers have been replaced with new identifiers. The reciprocal BLAST analysis yielded 73,023 RBHs for the *Synechococcus* queries and 5,410 RBHs for the *Synechocystis* queries. In comparison, the analysis by Schmelling et al., 2017 reported 24,027 RBHs for *Synechococcus* and 1,576 RBHs for *Synechocystis*. However, among all RBHs identified in this study, only 251 *Synechocystis* RBHs were not included in the previous analysis, while 4,907 *Synechococcus* RBHs were missed. Of the 251 *Synechocystis* RBHs, 28 have been removed from NCBI, 196 have been suppressed by NCBI, and 25 sequences do not reside in the database, leaving two sequences not detected in this analysis. Those two sequences can be found in the results of the forward BLAST, though. Of the 4,907 *Synechococcus* RBHs, 424 have been removed from NCBI, 2,548 have been suppressed by NCBI, and 327 sequences do not reside in the database, leaving 1,608 sequences not detected in this analysis. 118 of those sequences have been detected within the forward BLAST, while 1,490 sequences do not reside in the forward BLAST table.

The circadian clock proteins KaiA, Pex, LdpA, and CdpA have been exclusively identified in *Cyanobacteria* [37]. In the analysis conducted using CATHI, hits are observed outside the Cyanobacteria phylum for Pex and LdpA. Pex functions as a transcriptional repressor of kaiA in *Synechococcus elongatus* PCC 7942, featuring a relatively concise sequence length of 126 amino acids. For Pex, the CATHI pipeline identified 18 RBHs from 16 organisms beyond the *Cyanobacteria* phylum, displaying a mean bitscore value of 52 (Fig. 5). Moreover, none of those organisms possess RBHs of the central cyanobacterial circadian clock genes KaiABC. These results support the conclusion that these RBHs serve as indicators of distant homologous sequences, providing a compelling illustration of the necessity for user-driven post-processing — an aspect effortlessly facilitated through the interactive capabilities offered by CATHI. The phylogenetic tree derived from the analysis of those 18 non-cyanobacteria RBHs of Pex, in conjunction with a curated set of cyanobacterial Pex sequences, effectively illuminates the more distant relationship that exists among these sequences (Fig. 3). Notably, the redox-sensing protein LdpA demonstrates 268 non-cyanobacteria RBHs, characterized by a bitscore mean value of 218, signifying close homologous relationships. An intriguing pattern emerges within the 268 non-cyanobacteria hits for LdpA. A total of 248 hits are found within the *Streptophyta* phylum, originating from a diverse array of 68 distinct genera (88.46% of all *Streptophyta* possess RBHs of LdpA). Similar to the study conducted by Schmelling et al., 2017, RBHs of KaiA have been exclusively detected within the *Cyanobacteria* phylum in CATHI. Furthermore, an analysis of the KaiA protein sequence’s length distribution unveiled the same three distinct subtypes of KaiA (Fig. 7). Consistent with the analysis by Schmelling et al., 2017, the majority of KaiA RBHs exhibit a sequence length of approximately 300 amino acids, while the remaining RBHs vary between 100 and 200 amino acids in length. Our benchmarking analysis focused on the robustness and reproducibility of results when contrasted with previous analyses. While minor differences surfaced due to updates within NCBI’s database, these deviations did not impede the overarching tendencies (Fig. 7, Fig. S1). Notably, we noticed a large consistency within the analysis results even as the dataset increased by a factor of five compared to the previous study. This resilience underscores the enduring reliability of the methodology, firmly affirming its capability to withstand augmented data volumes without compromising the integrity and consistency of results.

**Figure 7.**
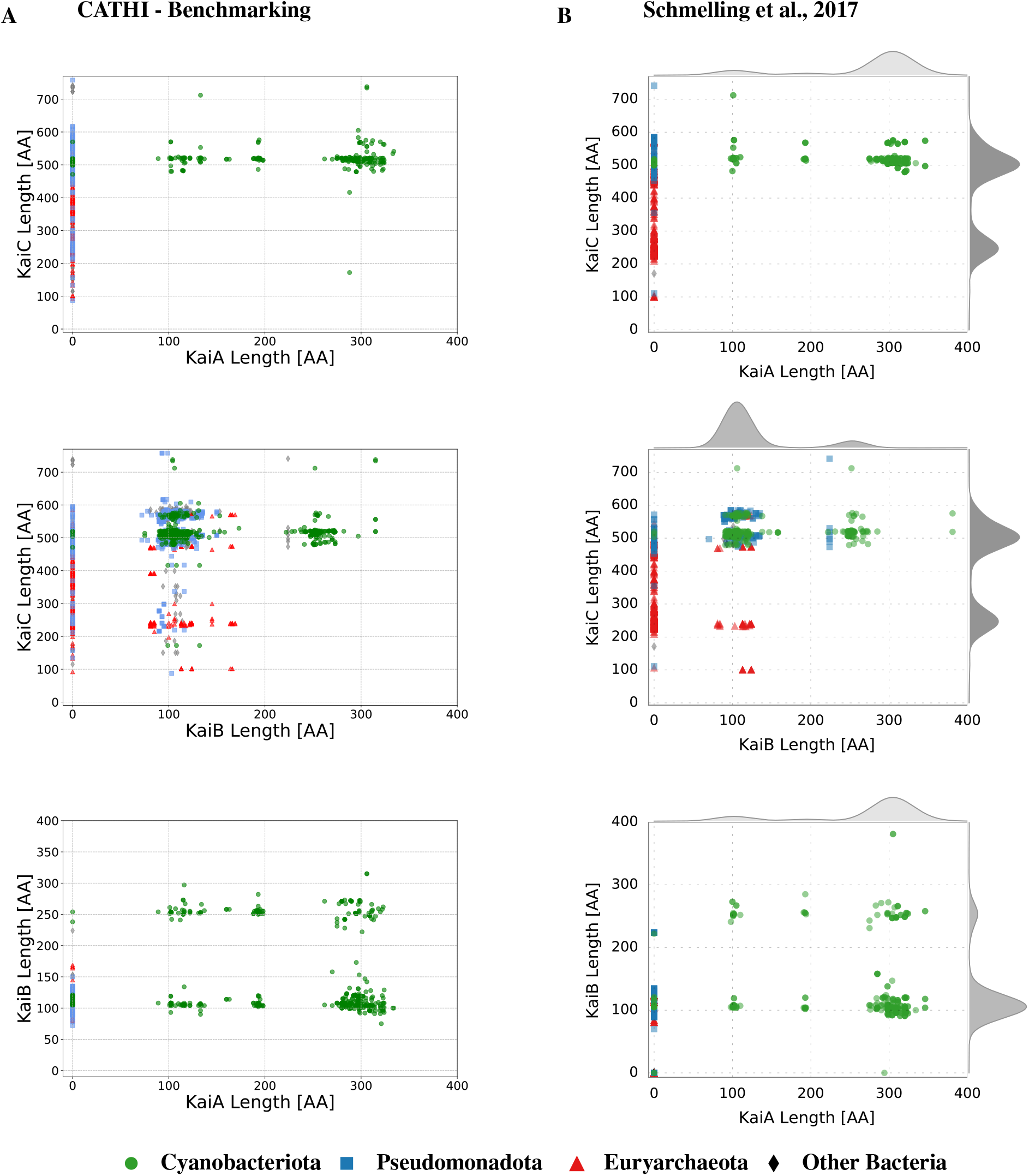
Comparison between the benchmarking analysis and Schmelling et al., 2017 of the sequence length distribution for the core circadian clock proteins KaiA, KaiB, and KaiC. This visual depiction offers an insightful comparison of sequence lengths for two of the three core circadian clock proteins using scatter plots. Four distinct organismic groups — *Cyanobacteriota, Pseudomonadota, Euryarchaeota*, and other bacteria — are the focal points of analysis. The overlay of the two datasets serves to highlight overarching patterns, underscoring the methodology’s robustness in elucidating sequence length dynamics across different organismic groups. Notably, this comparison not only reaffirms established trends but also emphasizes the enduring reliability of the analytical framework. This analysis showcases the consistency and robustness of the methodology and its capacity to yield insights that withstand scrutiny and validation. A) Results of the benchmarking analysis. B) Previous results from the reciprocal BLAST analysis of [37].

## 3 Methods

During the development of the application, all changes were consistently tracked with the version control system git. The code, its developmental history, and additional material can be found under the following link: https://github.com/Kanomble/celery_blast.

### 3.1 Docker-Based Software Deployment and Container Orchestration

The Docker container virtualization system was utilized to package the underlying software components into distinct images. Based on these images, seven containers were established, each designated for distinct tasks within CATHI. The images utilized for container creation are available on DockerHub under the following link: https://hub.docker.com/repositories/kanomble.

The base image of CATHI is constructed upon the Ubuntu Focal distribution and includes essential components such as the Django web framework (Version 4.2.4, employed for constructing the web server), Miniconda (Version 4.9.2, for Python package administration), the BLAST+ (Version 2.11.0) and the E-Direct software suite (Version 20.3.20230829) from NCBI, Snakemake workflow engine (Version 7.25.0), MAFFT (Version 7.453), FastTree2 (Version 2.1.11-1), and other critical software packages for running the web server and executing the pipeline. A comprehensive table of all utilized software tools and Python packages can be found in Supplementary Table S2. Four Docker containers are derived from this foundational base image:

- The **web** container orchestrates the startup of the Django application (including database migrations) and the Python WSGI server, Gunicorn (Version 20.1.0).
- The **celery_worker** container utilizes a local celery installation (Version 5.3.1), which handles long-running background tasks, such as the Snakemake pipeline.
- The **flower** container is a monitoring interface for celery tasks and workers, utilizing the Flower (Version 2.0.1) web server.
- The **jupyter_notebook** container launches the web-based interactive computing platform Jupyter (Version based on IPython 8.12.0).

The foundational database of the web server is encapsulated within a PostgreSQL container. During the initial startup of the web container, database tables, and base models are written into the database, a process facilitated through the use of the Django web framework’s makemigration and migrate commands. These programs translate Django’s model classes into SQL tables (known as Object Relational Mapping, ORM).

The rabbitmq container is dedicated to launching the message broker RabbitMQ (Version 3.9.4), which takes user-generated task instructions and conveys them to the celery framework.

CATHI employs Nginx (Version 1.21.6) as a reverse proxy, directing client requests to a Gunicorn backend server and relaying the responses to clients. This pivotal role distributes incoming traffic, enhancing the system’s overall performance. The initiation of Nginx is executed through the dedicated Nginx Docker container.

To ensure proper container initialization, a specific startup sequence is vital. The web container necessitates the database container’s availability; similarly, the celery container relies on the functioning rabbitmq container. The wait-for script (Version 2.1.0), reliant on netcat (Version 1.206-1ubuntu1), is employed to manage this sequence. Written in BASH, this script accepts a port number as input and continuously checks for the presence of an application listening on that port. Once a port is occupied, the script ceases its blocking function, allowing the container to initiate its operations. Once launched, users can access the application via common browser applications using this address: http://127.0.0.1:1337/blast_project/.

### 3.2 Django Web Framework

For an efficient, clean, and pragmatic design, the high-level Python web framework Django (Version 4.2.4) was used for the development of this web application. Django, a popular high-level web framework written in Python, accelerates web application development through its Model-View-Controller (MVC) architecture, integrated ORM, and dynamic template engine. Utilizing Django, the integration of additional Django-based applications is achievable by including the respective software packages within the INSTALLED_APPS environment variable, as specified in the settings.py file of the project. The following third-party applications have been incorporated into CATHI: django_extensions (Version 3.2.3), django_celery_results (Version 2.5.1), and celery_progress (Version 0.3). The django_extensions application facilitates the execution of the startup.py script, which introduces database models into the PostgreSQL (Version 13.4) database and procures the taxonomy database from NCBI during the initial launch of the web container. In parallel, the django_celery_results application extends functionality by enabling the storage of celery task outcomes through the utilization of the Django ORM framework.

### 3.3 Web Development Tools

CATHI uses Django’s built-in template language for HTML documents and employs cutting-edge web technologies to create interactive and visually appealing websites. The use of JavaScript and AJAX guarantees seamless web interactivity, complemented by Bootstrap 5.0 for bolstering aesthetics through predefined Cascading Style Sheet (CSS) components. Additionally, the HTML table plugin for JavaScript, DataTables optimizes the presentation and manipulation of HTML tables, resulting in a comprehensive toolkit specifically designed to create dynamic and user-friendly web experiences. All third-party web libraries are integrated via their respective content delivery networks (CDNs). Custom CSS and JavaScript code are located within the static directory of CATHI.

### 3.4 Project Creation

Project creation is facilitated by Django’s built-in form validation. User input is validated before a project is manifested into the PostgreSQL database. There are certain input requirements for the reciprocal BLAST projects. The SOIs have to come from only one organism per project and they have to reside in the backward database. The best option is to upload the genome from which the user has obtained the sequences with CATHI’s database creation module. The input is validated by Django’s internal form validation functionality and accurate failure messages are displayed if form validation fails. In addition to the SOIs and the species name of the target organism, users have to enter a project title, which has to be unique, no other project should use this title. In addition, users can adjust BLAST settings (for forward and backward BLAST separately), apply a bitscore filter and limit the maximum number of RBHs used to create a phylogeny, as well as change settings of the post-processing programs trimAl [45] and MView [46].

### 3.5 Pipeline Execution with the Snakemake Workflow Engine

The CATHI pipeline process leverages the Snakemake [51] workflow engine to streamline the entire process of RBH analysis. This integration not only increases the reproducibility of results but also reduces the burden on the user by automating many of the necessary steps in the detection of putative homologs. The developed Snakemake pipelines integrate a diverse array of algorithms and software tools. Three Snakefiles have been developed: one for the core reciprocal BLAST pipeline, another for local BLAST operations, and a third for conducting remote BLASTs on NCBI servers. Alongside supplementary scripts, these Snakefiles reside within dedicated static subdirectories.

Snakemake is executed within a celery task encapsulated in a Python subprocess.Popen call. The core Snakemake pipeline for detecting RBHs comprises the following steps:

1. Forward BLAST
2. Backward BLAST preparation
3. Backward BLAST
4. Extraction of RBHs (this is done via pandas merging tools)
5. Post-processing of RBHs (inference of taxonomic information, statistics, HTML and CSV tables, basic result plots)
6. Extraction of RBH-sequences separated by query sequences
7. MSA of each set of RBHs with MAFFT
8. Phylogenetic inference of each set of RBHs with FastTree2
9. Post-processing of the phylogenetic tree and the MSA with shiptv, trimAI and MView
10. CDD domain search of target sequences

### 3.6 Database Creation

CATHI enables the creation of local BLAST databases based on publicly available or uploaded protein sequences. For the publicly available sequences, CATHI utilizes the RefSeq and GenBank assembly summary files, which inherit FTP paths to the corresponding protein assemblies. The assembly summary files can be filtered at two different levels. The first level includes the completeness status of the dedicated assembly, which corresponds to the genome coverage and completeness of the genome assembly. This is summarized by an assembly level of the corresponding protein genome file. The second filtering step involves organism-specific filtering. For each organism and clade of the taxonomic classification (e.g., for genus, family, or order), a taxonomic node is stored in the taxonomy database [55]. These taxonomic nodes can be used to filter the assembly summary file to create BLAST databases containing only organisms with the provided taxonomic nodes. Taxonomic node files can be uploaded directly as part of the database formatting process or created beforehand by using CATHI, which provides a dashboard for translating higher taxonomic nodes to the underlying species-specific nodes. This translation procedure is realized by executing the Perl script get_species_taxids.pl provided by the BLAST+ command line tool.

Sometimes, it is necessary to work with unpublished genomes or sequences. CATHI offers two options for uploading your own protein FASTA files, which are then formatted into BLAST databases. The user can upload a concatenated protein FASTA file consisting of multiple genomes. This approach requires additional metadata; the assembly identifier (combination of characters or numbers), organism names (with valid taxonomic nodes), assembly levels (one out of four; “Contig,” “Scaffold,” “Chromosome,” or “Complete” Genome), and a file that contains a mapping of the provided sequence identifiers to the relevant taxonomic nodes (taxmap file). This approach is error-prone and may lead to problems for inexperienced users. Therefore, a second approach was developed, which allows the uploading of multiple single protein FASTA files in combination with valid species names. Uploaded files are then used to build BLAST databases with the makeblastdb program.

Prior to the formatting procedure using makeblastdb, protein FASTA files undergo parsing, and the header of each sequence is slightly modified to incorporate the genome assembly name, thus assigning a unique identifier to each sequence. This approach mitigates the issue of multiple identical sequence identifiers within the database, which could otherwise trigger an error during the makeblastdb database formatting process. BLAST databases are divided into chunks, with each chunk comprising 500 genome files. These individual database chunks are subsequently compiled into a single .pal alias file, consolidating all database chunks into a unified BLAST database. This method prevents excessive RAM consumption for extensive databases and facilitates subsequent database updates.

### 3.7 BLAST and Identification of Reciprocal Best Hits

The forward and backward BLAST (Fig. 1) analysis of the user-provided SOI is performed using the blastp program within the BLAST+ software suite (Version 2.11.0) [15]. The default settings for the forward BLAST are as follows: e_value=0.001, word_size=3, num_alignments=10.000, max_hsps=500. The default settings for the backward BLAST are similar to the forward BLAST settings, except for the num_alignment option, which is set to num_alignments=1. Default settings can be customized by the user. Other tools within the BLAST+ software suite utilized in CATHI include the makeblastdb program, employed for formatting protein BLAST databases; the blastdbcmd program, which facilitates the retrieval of protein sequences identified during the initial BLAST analysis; and the rpsblast program, which is used for the inference of CDDs among the RBHs. RBH inference is executed using the Python library Pandas, which involves a comparison between the outcomes of forward and backward BLAST analyses. Additionally, the BioPython library is harnessed to deduce the underlying taxonomic details of the organisms of the resulting RBH sequences, which are then documented in a CSV file. All tables generated are conveniently accessible within CATHI, facilitated by integrating the client-side JavaScript library DataTables (see Methods - Web Development Tools).

### 3.8 Post-Processing Software

Python’s Biopython (Version 1.78) [44] library is used to extract taxonomic information from database entries. More-over, Biopython is used to slice protein GenBank files acquired through synteny analysis conducted by CATHI. These segmented GenBank files are subsequently used as input for the clinker tool (Version 0.0.27). Matplotlib (Version 3.7.2) serves to generate fundamental result graphs, pandas (Version 1.2.4) facilitates the manipulation of tabular data and the inference of RBHs, while scipy (Version 1.10.1) and scikit-learn (Version 1.3.0) are utilized for Principal Component Analysis (PCA) within CATHI’s CDD detection module. Bokeh (Version 2.4.3) is used to create interactive visualizations (based on the database and the RBH result tables or the CDD result table). Third-party tools MAFFT (Version 7.453) and FastTree2 (Version 2.1.11-1) are employed for MSAs and the inference of corresponding phylogenetic trees. TrimAl (Version v1.4.rev22) is used to trim the MSA by removing long gaps and uninformative segments from the alignment. MView (Version 1.67) and shiptv (Version 0.4.1) transform the output of these tools to generate interactive HTML documents for MSAs and phylogenies. A comprehensive table of all employed software tools and Python packages can be found in Supplementary Table S2.

## 4 Discussion

Identifying closely and distantly related homologous sequences is an essential task in the field of comparative genomics [1, 2, 56]. Typically, orthologous and paralogous gene relationships are differentiated by evaluating and contrasting sequence similarities and their distribution within phylogenetic contexts. However, diverse biological questions may necessitate distinct computational approaches, leading to the development of various programs (Table 1) for homology inference [6, 9]. Furthermore, over the past decades, biological databases with collections of orthologous sequences have emerged, expediting the swift identification of orthologous relationships within established sequences [24, 57, 25, 58, 27, 26]. The ever-increasing biological data demands flexible approaches to disentangle homologous relationships among genes, within newly sequenced genomes, which are not part of these databases. CATHI is a platform for the interactive exploration of the output of sequence similarity searches in custom databases that can be easily created via dedicated web-interfaces. The additional post-processing modules, such as phylogeny, synteny, and inference of conserved domains among identified homologous sequences, make CATHI a highly flexible tool that enables rapid and interactive analysis of results without the need for additional bioinformatics tools. In this chapter, we discuss CATHI’s usability and highlight advantages and disadvantages in comparison to other, similar computational resources.

### 4.1 Usability

There are versatile tools for inferring homologous sequences (Table 1). While tools like PARIGA [59] and orFin [10] are no longer accessible, other tools like morFeus, JustOrthologs, and orthoFinder offer different strategies for the identification of orthologs. The later tools are either accessible through the command line or hosted on dedicated web-servers. In contrast, CATHI introduces a unique paradigm. It presents an innovative approach by providing the flexibility of both local and server site installations, coupled with an interactive interface accessible via standard web browsers. Most tools provide comprehensive tables of results listing the putative orthologous sequences identified in different organisms (Table 1). However, a thorough analysis of these results often requires additional software tools and further post-processing steps. Tasks such as MSAs and phylogenetic tree inference frequently entail programming expertise, posing challenges for many biologists attempting to address these complexities.

Drawing from our experience, a significant portion of biologists exhibit concerns regarding coding prerequisites. Consequently, a notable demand exists for software tools that provide graphical user interfaces, facilitating result analysis through user-friendly cursor interactions and drag-and-drop functions. Furthermore, manual intervention and examination are often necessary to analyze homologous sequences, such as distinguishing distantly from closely related homologs. Hence, an intuitive interface that streamlines result analysis and integrates crucial post-processing steps is essential for homologous sequence analysis. CATHI offers such interfaces by using e.g. the capabilities of the interactive plotting library Bokeh. Users can employ these interfaces to filter results based on taxonomy and BLAST statistics, as well as interactively select RBHs for various subsequent analyses (Fig. 5). Examining the RBHs of the *Synechococcus* Pex protein within the *Bacillota* and *Bacteriodota* phylum, a total of 18 RBHs from 16 organisms of the Pex protein have been found, characterized by relatively modest bitscores (mean of 52). While these identified RBHs might represent homologous sequences, their lower bitscores and reduced sequence identity could doubt their status as orthologous sequences. This underscores the need for user intervention and careful consideration through evaluating outcomes. CATHI enables such manual post-processing with a comprehensive graphical display that allows effortless filtering and editing of results (Fig. 5). In addition, CATHI uses standard bioinformatics algorithms whose software versions are embedded in the Docker image. This arrangement increases the stability of the pipeline in executing consistently, even if software versions change, and if a database update is required, users can use CATHI’s database creation module and rerun the pipeline, ensuring robust and reproducible gene relationship detection. Nonetheless, bioinformatics software tools designed for orthologous sequence detection can assist in estimating the potential presence of homologous sequences. However, to validate a closely related homolog as a *bona fide* ortholog, it is essential to conduct wet lab experiments focusing on protein functionality.

### 4.2 Strength and Drawbacks in Homolog Identification in CATHI

The method of deducing RBHs as potential homologs may not invariably yield accurate results [60], particularly in cases involving ancient gene duplications followed by selective paralog loss [61], which can partly be resolved by synteny analyses. Consequently, the extant paralog might erroneously be categorized as an ortholog. In a broader context, if the organisms of interest harbor only a single homolog of the query sequence, and a prospective organism possesses a solitary copy of a homologous gene, that single copy could manifest as an RBH in the results, even if it represents a closely related homolog rather than a genuine ortholog. There are other tools that implement more sophisticated approaches to detect the orthologous and paralogous relationship among genes (Table 1) [60]. In the upcoming section, we will contrast OrthoFinder [12], one of the most advanced tools for ortholog detection, with CATHI. This will allow us to highlight the strengths and limitations of these two tools, emphasizing the key distinctions that prompted the development of CATHI.

OrthoFinder assigns genes to orthogroups using an orthogroup inference algorithm, infers rooted trees using the STAG and STRIDE programs [31, 32], and then applies a hybrid algorithm based on the species-overlap method [33] and the duplication-loss-coalescent model [34] to identify orthologs and paralogs. In contrast to the RBH approach, this method leads to sophisticated ortholog and paralog assignments. However, the output of OrthoFinder are text files that need to be analyzed in additional post-processing steps outside of OrthoFinder. Furthermore, OrthoFinder’s search space is constrained by the number of input FASTA files, and it focuses on whole genomes rather than individual sequences. It does not infer orthologs in comprehensive databases, which limits the potential to unveil certain features of the SOIs when searching for homologs in extensive databases.

If the number of gene duplications is relatively high, such as in angiosperms [62], the RBH approach may not detect all orthologs, only those with the best alignment scores are reported [63]. In this case, OrthoFinder and other tools that make use of calculating orthogroups yield more accurate results compared to the RBH approach implemented in CATHI [64, 60]. However, at least for prokaryotic genomes the RBH method can still serve as a strong indication for gene orthology [65]. In addition, CATHI’s custom databases can expand the search scope for identifying novel orthologous or paralogous genes, particularly by allowing rapid, visual filtering of results obtained from the analysis, whereas OrthoFinder is limited to the number of FASTA files entered. CATHI also provides options for simple one-way BLAST analyses that can help identify extended duplication events when there are multiple homologous sequences in an organism with relatively high statistical significance (e.g., bitscores > 50 or e-values < 0.001).

However, the exact classification of genes as orthologs or paralogs is difficult to implement even for tools like OrthoFinder. To safely predict orthology, we would need a complete history of all ancestral genomes. To date, our genomic information is still insufficient and might be forever as genetic information throughout the history of life has been irrevocably lost. Thus, predictions will only yield reliable results to a certain extent. Even though it is an interesting and important question, for most laboratory biologists those distinctions are not their main focus. They rather use homology searches to better understand their protein/gene of interest before examining it in more detail in the laboratory. Here, indications about potential functions, involvement in metabolic or regulatory pathways, or location on the genome within clusters or in proximity to other well characterized genes is far more useful. In general, with CATHI it is possible for users inexperienced with command line tools or programming languages to gain insights about a gene of unknown function by identifying homologs and analyzing the synteny, phylogeny, and domain structure in an automated workflow, while exploring and filtering the data in an interactive interface. That information will then guide laboratory experiments to correctly describe the function of the gene.

CATHI utilizes some generalizations that introduce certain limitations, such as using the blastp program for conducting the sequence similarity search algorithm, thus, restricting the acceptable query sequences to proteins. Furthermore, it uses a reciprocal BLAST to search for close homologous sequences. While other tools may be more accurate at identifying orthologs [63], CATHI provides intuitive results analysis suitable for both experienced and less experienced users. The automated pipeline creates a generalized workflow for homolog identification. Nevertheless, CATHI may not be suitable for all SOIs due to the underlying reciprocal BLAST technique. However, it provides a great starting point for most projects, thereby extending the accessibility of homologous sequence analysis to a broader spectrum of users. Through interactive filtering, more experienced users can reduce the initial BLAST cut-offs to allow for a broader homology search and later filter those results using the extensive options in the post-processing module.

### 4.3 Database Content and Customization

Certain tools possess a restricted search space attributed to static databases (e.g., InParanoiDB9 or COG) and/or the necessity for pre-formatted databases (e.g., OrthoInspector, morFeus or JustOrthologs), which subsequently remain “static” in a manner that necessitates users to download and format each database in advance.

Due to the rapid growth of biological data, a flexible tool is needed that can automate rapid configuration of biological databases. CATHI provides the capability to generate customized BLAST databases utilizing publicly accessible and locally available protein sequences. Upon completing the database creation process, a dedicated webpage exhibits a table showcasing the database’s contents. This comprehensive table enumerates all organisms and genome entries encompassed within the database, facilitating researchers in obtaining an overview of its contents. Furthermore, databases can be customized based on criteria such as completeness level, taxonomy, and the option to download either from RefSeq or GenBank, consequently facilitating more nuanced scrutiny of sequence similarity searches, which is vital in detecting specific homologous sequences. Once the database is downloaded, its entries remain unaltered, simplifying the monitoring of modifications in publicly accessible databases. This feature serves to alleviate potential confusion stemming from omitted or deleted sequence entries and to overcome limitations within the provided search scope.

### 4.4 Modular Expandability

The use of Docker, Django, and Snakemake provides flexibility in extending the pipeline to include additional functionality or analytics. The modular nature of these technologies allows for seamless integration of new functionality and increases the adaptability and future-proofing of the pipeline. Typically, bioinformatic tools are bundled into software packages that are executed on the UNIX command line (e.g. OrthoFinder). These packages often work independently and perform the desired task with certain inputs provided by the user. However, customizing these programs to meet specific requirements demands users to possess experience with the underlying code of these tools. On the other hand, extending CATHI’s Snakemake pipeline is relatively easy for sophisticated users, giving rise to the opportunity to create novel bioinformatic workflows targeting specific research questions. Snakemake defines a workflow using a directed-acyclic-graph (DAG) builded through a set of rules whose execution is based on certain input files and some additional parameters. These rules generate output files, which in turn can serve as input files for other rules. Users can extend the provided Snakemake workflow by adding additional rules that can make use of the infrastructure generated by CATHI.

## 5 Outlook

In its current version, CATHI provides a simple entry point into comparative genomic analysis for biologists. However, thanks to its modular structure, the platform can easily be extended. The platform’s homolog search module could evolve to allow users to opt for different search tools like BLAST or DIAMOND [16], improving the efficiency of homology searches. The integration of advanced techniques such as Markov clustering, hidden Markov models (HMMs) of sequence alignments that can be used to detect distant homologs [66], and bootstrapping into the homolog search process has the potential to enhance the precision of homolog identification, advancing the accuracy of comparative genomics analyses. Furthermore, the incorporation of iterative homology searches introduces a refinement process. Users can initiate searches, refine them through filtering, and adapt parameters based on evolving insights, ensuring a dynamic and adaptive analytical workflow. Future iterations of CATHI can encompass sophisticated co-occurrence analyses of RBHs, unveiling intricate patterns of protein relationships across diverse organisms. In addition, the inclusion of additional tools such as OrthoFinder and TreeViewer holds the potential to enrich the analytical spectrum, broadening the toolbox available to researchers. Last, the untapped potential of emerging AI tools and capabilities holds the promise to further augment this platform’s functionalities, potentially unearthing novel and unforeseen avenues for analytical insights. Incorporating these enhancements will bolster the platform’s versatility, ensuring its alignment with the evolving landscape of bioinformatics tools and methodologies.

