## Supplementary Data for "CATHI: An interactive platform for comparative genomics and homolog identification"

Table S1: **Protein input queries for the reciprocal BLAST benchmarking analysis.** Cyanobacterial circadian clock proteins were selected from *Synechococcus elongatus* PCC 7942 and *Synechocystis* sp. PCC 6803, based on the previous analysis by Schmelling et al., 2017 [1].

| Organism Name | Protein Name | RefSeq ID |
|---|---|---|
| *Synechococcus elongatus* PCC 7942 | KaiA | WP_011377921.1 |
| | KaiB | WP_011242647.1 |
| | KaiC | WP_011242648.1 |
| | Pex | WP_011377679.1 |
| | LdpA | WP_011377652.1 |
| | NhtA | WP_011378346.1 |
| | PrkE | WP_011243235.1 |
| | CdpA | WP_011378107.1 |
| | CikA | WP_011243194.1 |
| | SasA | WP_011378322.1 |
| | LabA | WP_011244514.1 |
| | LalA | WP_011242719.1 |
| | Crm | WP_011243720.1 |
| | RpaA | WP_011377437.1 |
| | RpaB | WP_011378039.1 |
| | CpmA | WP_011377895.1 |
| *Synechocystis* sp. PCC 6803 | KaiB1 | WP_010874242.1 |
| | KaiC1 | WP_010874243.1 |
| | KaiB2 | WP_010872548.1 |
| | KaiC2 | WP_010872549.1 |
| | KaiB3 | WP_010874242.1 |
| | KaiC3 | WP_010873229.1 |

Table S2: **Selection of CATHIs software tools and version numbers.**

| **Tool** | **Version** |
|---|---|
| BLAST+ | 2.11.0 |
| EDirect | 20.3.20230829 |
| MAFFT | 7.453 |
| FastTree2 | 2.1.11-1 |
| PostgreSQL | 13.4 |
| RabbitMQ | 3.9.4 |
| nginx | 1.21.6 |
| MView | 1.67 |
| trimAl | v1.4.rev22 |
| netcat | 1.206-1ubuntu1 |
| wait-for | 2.1.0 |
| clinker | 0.0.27 |
| shiptv | 0.4.1 |
| Snakemake | 7.25.0 |
| bokeh | 2.4.3 |
| scikit-learn | 1.3.0 |
| scipy | 1.10.1 |
| pandas | 1.2.4 |
| matplotlib | 3.7.2 |
| biopython | 1.78 |
| IPython | 8.12.0 |
| Miniconda | 4.9.2 |
| conda | 23.7.3 |
| Django | 4.2.4 |
| django-celery-results | 2.5.1 |
| django-extensions | 3.2.3 |
| flower | 2.0.1 |
| gunicorn | 20.1.0 |
| celery | 5.3.1 |
| celery-progress | 0.3 |

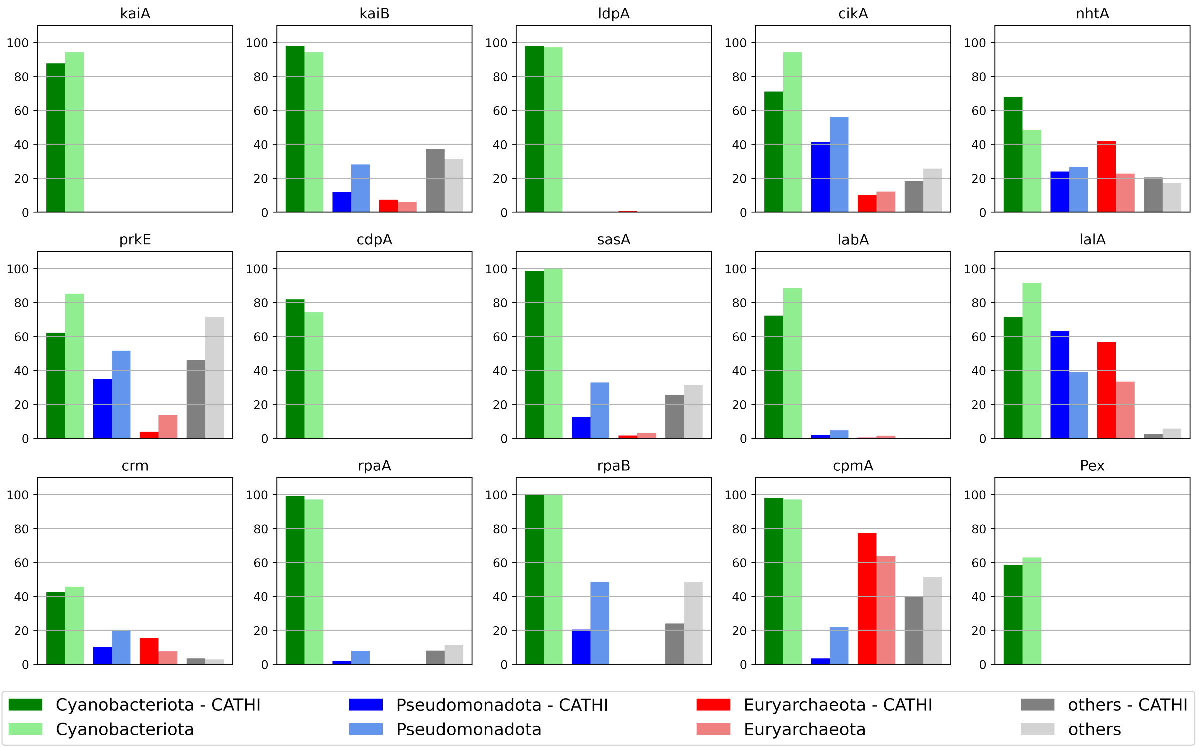


Figure S1: **Comparison between CATHI and Schmelling et al., 2017 of the abundance within the RBHs of the circadian clock core-, input- and output-factors of organisms that harbor a KaiC homolog.** For each circadian clock protein, the abundance value is calculated as the ratio of the number of RBHs for the circadian clock protein divided by the number of organisms with RBHs for KaiC. While there are some differences between the studies, the overall trend remains similar. Darker colors refer to the results of this benchmark analysis, while lighter colors refer to the results of the initial study by Schmelling et al., 2017 [1]. Green: *Cyanobacteriota*; Blue: *Pseudomonadota*; Red: *Euryarchaeota*; Other Bacteria.
